## Supplementary figures and images for "E3 ubiquitin ligase Deltex facilitates the expansion of Wingless gradient and antagonizes Wingless signaling through a conserved mechanism of transcriptional effector Armadillo/β-catenin degradation"

### Supplemental Figure 1

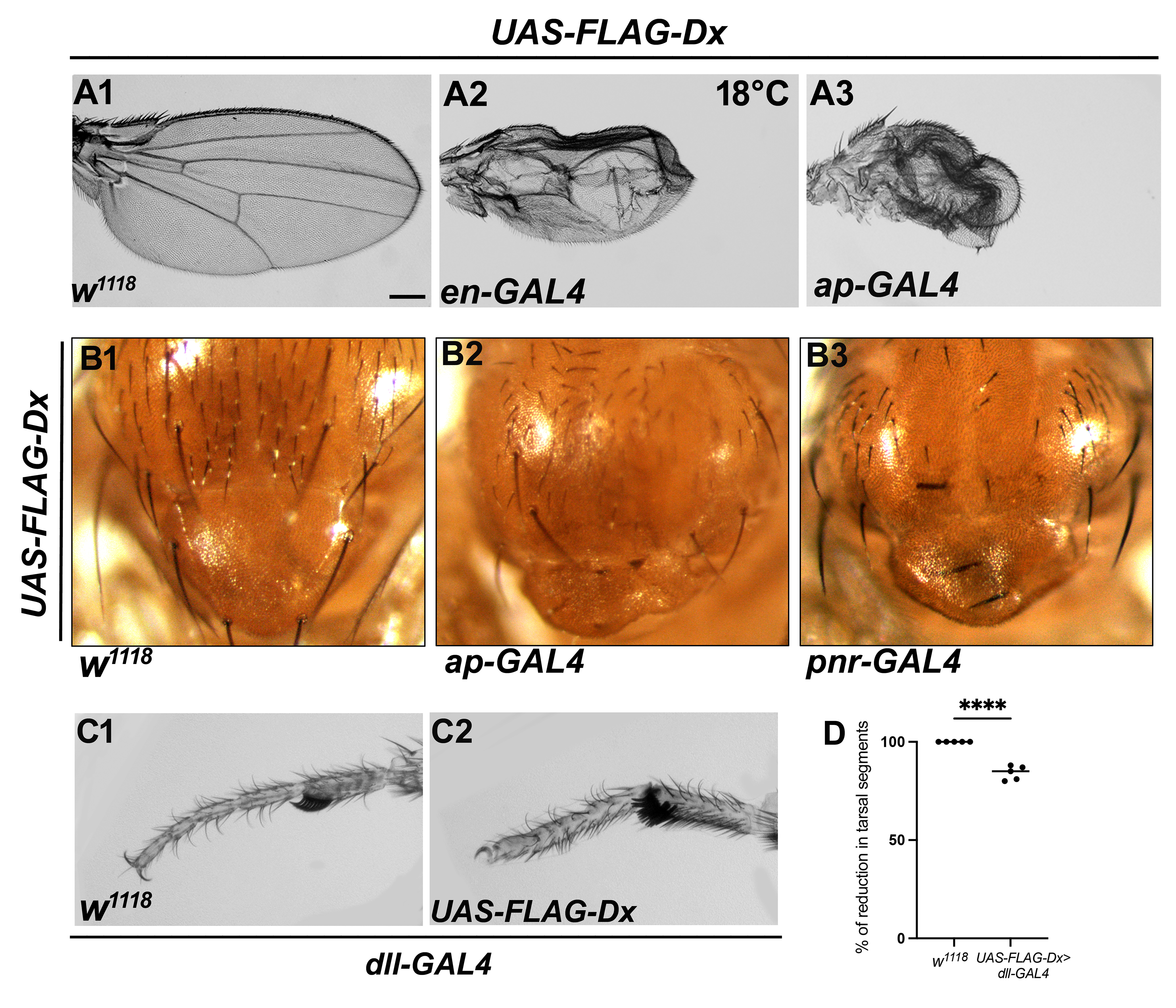

### Supplemental Figure 2

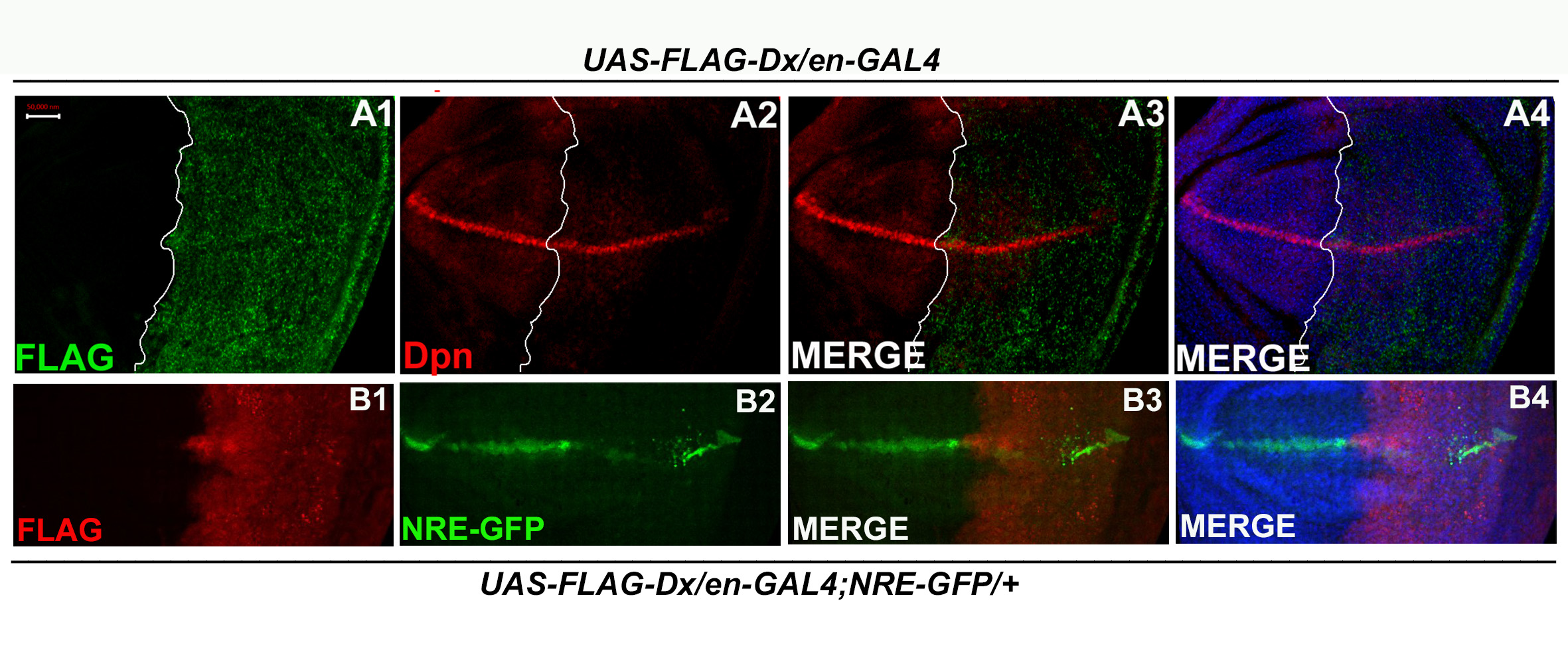

### Supplemental Figure 3

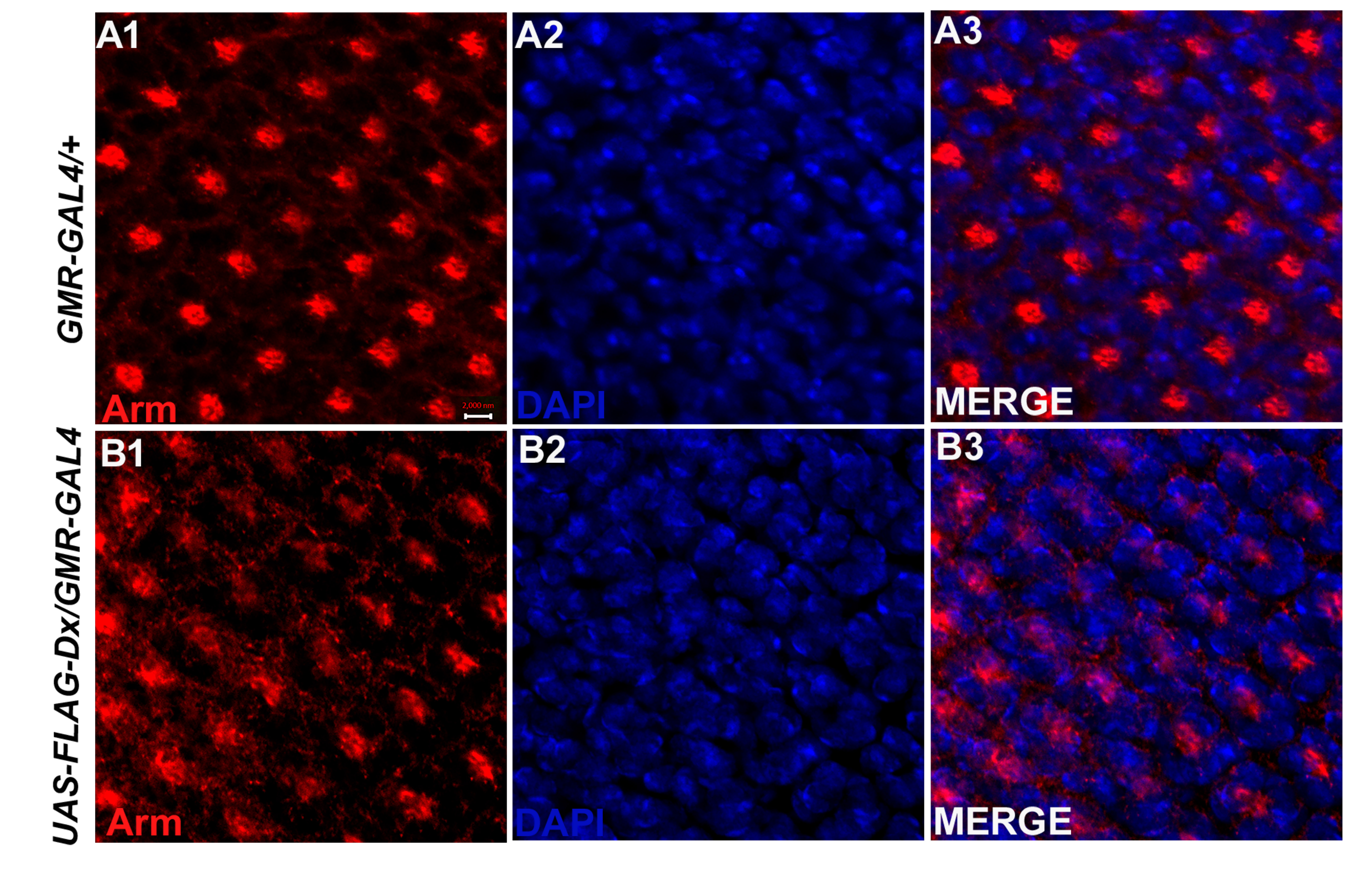

### Supplemental Figure 4

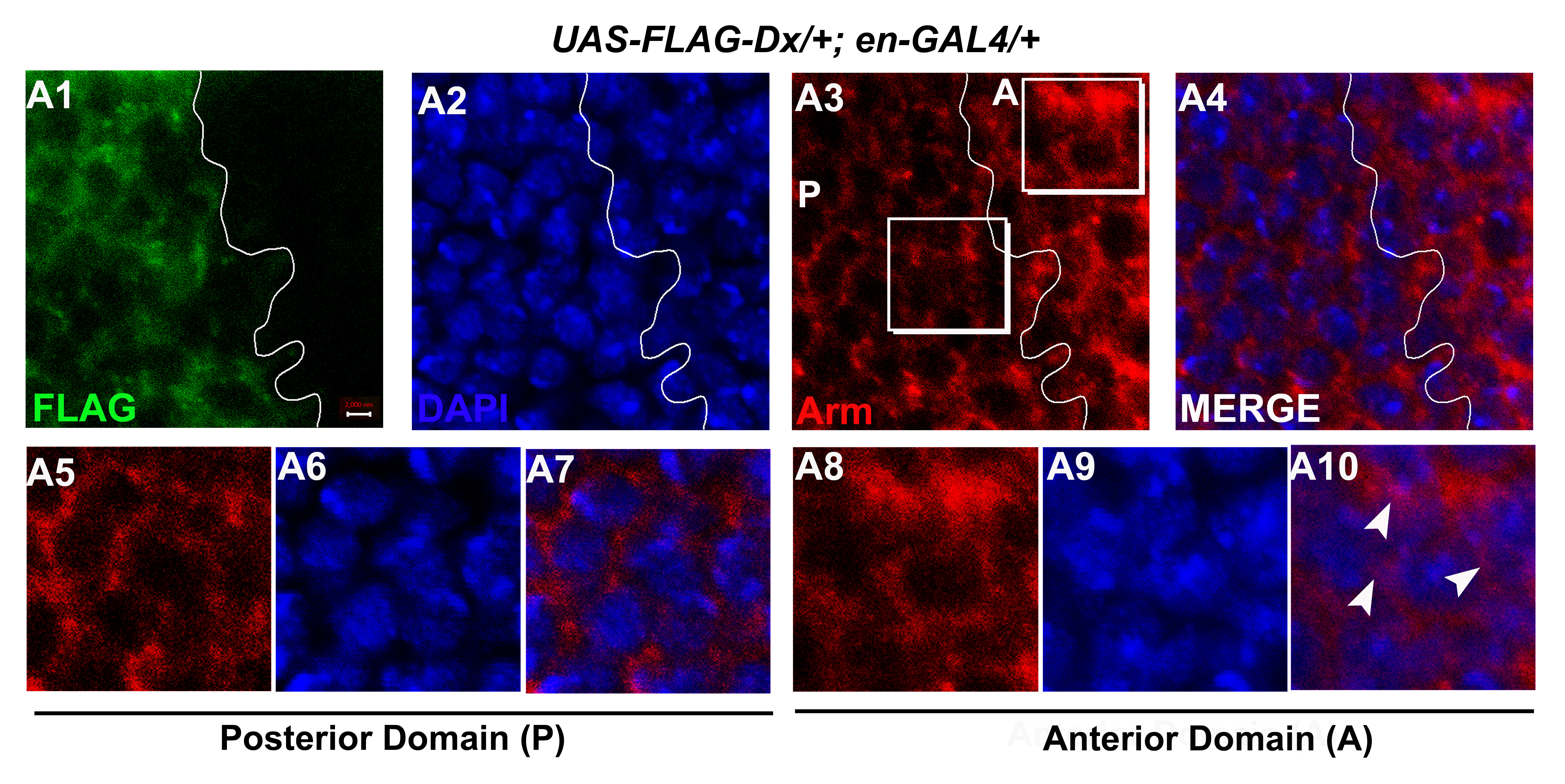
